# *In vitro* reduction of *Plasmodium falciparum* gametocytes by *Artemisia spp.* tea infusions

**DOI:** 10.1101/2020.07.29.227983

**Authors:** Danielle Snider, Pamela J. Weathers

## Abstract

In this study, we showed *in vitro* evidence that supports the efficacy of *A. annua* and *A. afra* tea infusions used in a 2015 clinical trial not only for clearing asexual *Plasmodium falciparum* parasites, but also for eliminating sexual gametocytes. *P. falciparum* NF54 was grown *in vitro,* synchronized, and induced to form gametocytes using N-acetylglucosamine. Cultures during asexual, early, and late stage gametogenesis were treated with artemisinin, methylene blue, and *Artemisia annua* and *A. afra* tea infusions (5g DW/L) using cultivars that contained 0-283 μM artemisinin. Asexual parasitemia and gametocytemia were analyzed microscopically. Gametocyte morphology was also scored. Markers of early *(PfGEXP5)* and late stage *(Pfs25)* gametocyte gene expression were also measured using RT-qPCR. Both *A. annua* and *A. afra* tea infusions reduced gametocytemia *in vitro*, and the effect was mainly artemisinin dependent. Expression levels of both marker genes were reduced with the effect mainly attributed to artemisinin content of the our tested *Artemisia* cultivars. Tea infusions of both species also inhibited asexual parasitemia and although mainly artemisinin dependent, there was a weak antiparasitic effect from artemisinin-deficient *A. afra*. These results showed that *A. annua* and to a lesser extent, *A. afra,* inhibited parasitemia and gametogenesis *in vitro,* and results are consistent with earlier observed clinical results.

## Introduction

Malaria is a severe global health problem that disproportionately affects Africans and especially children under age 5. In 2018, there were 228 million cases of malaria worldwide, and 93% occurred in Africa (World Health Organization, 2019). The plant *Artemisia annua* L. has been used for > 2,000 years to treat fever, a characteristic of malaria. Artemisinin is the antimalarial sesquiterpene lactone isolated from the glandular trichomes of this plant. Artemisinin has poor solubility and low bioavailability, so it is no longer clinically used. Instead, it has been replaced with one of four semisynthetic derivatives used in combination with a partner drug to form artemisinin-combination therapies (ACTs), the current frontline global antimalarials (Gomes *et al.*, 2016). To achieve eradication of malaria, therapies must not only eliminate patient infections, but also prevent parasite transmission (The malERA Consultative Group on Drugs, 2011).

When the malaria parasite, *Plasmodium falciparum,* enters the human body via a mosquito bite, it first undergoes asexual development, followed later by its sexual stage (Phillips *et al.*, 2017). The asexual stage causes the severe clinical symptoms of the disease and, if left untreated, can result in death. The sexual stage, gametocytes, do not contribute to patient mortality, but rather are responsible for parasite transmission back to the mosquito to complete the full life cycle of the parasite (Phillips *et al.*, 2017). Gametocytes are also crucial therapeutic targets, because by eliminating them, the cycle of malaria can be broken.

Few currently used antimalarials are effective at eliminating both stages of the parasite. Most antimalarials target metabolically active parasite stages (Delves *et al.*, 2013). As gametocytes mature, however, their overall metabolic activity declines until they reach quiescence at maturity (Young *et al.*, 2005). While some antimalarials, including artemisinin derivatives, chloroquine, quinine, and atovaquone, have some activity against early stage gametocytes, only primaquine is clinically approved to kill late stage gametocytes (Baker, 2010; Duffy and Avery, 2013; Beri *et al.*, 2018). Primaquine, however, has some serious adverse effects, so safer gametocyte-targeted therapeutics are desirable (Sanofi-Aventis, 2017).

In 2015, a clinical trial compared the efficacy of a standard ACT, artesunate-amodiaquine (ASAQ), against tea infusions of *A. annua* and its cousin *Artemisia afra* (Munyangi et al. 2019). Not only did both *Artemisia* infusions eliminate asexual-stage parasites more effectively than ASAQ (88.8% for *A. afra,* 96.4% for *A. annua,* vs. 34.3% for ASAQ at day 28), but both tea infusions were also more effective in clearing microscopically measured gametocytes from the bloodstream (100% for both teas vs. 98% for ASAQ by day 14) (Munyangi *et al.*, 2019).

Based on those clinical results, it is important to better understand the gametocytocidal effects of the two *Artemisia* species by comparing how each tea infusion treatment affects gametocytes at different stages of their maturation. Gametocytes are significantly under-estimated when only measured microscopically, so qPCR analysis of developmental stage markers is important to establish the efficacy of any anti-gametocyte therapeutic (Bousema *et al.*, 2006). Here we used RT-qPCR to track the early and late stage markers, *PfGEXP5* and *Pfs25,* respectively, to further measure the efficacy of *Artemisia spp.* tea infusions *in vitro* against NF54 *P. falciparum* gametocytes.

## Materials and Methods

### Plant material and its preparation for testing

*Artemisia annua* L. cv. SAM (voucher MASS 317314) and *A. afra* Jacq. ex Willd. (SEN, voucher LG0019529; PAR, voucher LG0019528; LUX, voucher MNHNL2014/172) tea infusions were all prepared from dried plant material (leaves and small twigs) steeped in boiling water for 10 minutes to create a final concentration of 5 g/L. After cooling, the infusion was then successively filtered as follows: 1 mm sieve, 600 μm sieve, Whatman #1 filter paper, Millipore RW03 pre-filter, 0.45 μm type HA filter, and last a 0.22 μm filter to sterilize. Sterile infusion was aliquoted into 1.5 mL tubes and stored at −80°C. Artemisinin content of the filtered infusions was determined by gas chromatography mass spectroscopy (GCMS) as detailed in Martini et al. 2020 (Martini *et al.*, 2020). SAM *A. annua* and SEN *A. afra* tea infusions contained 283 and 0.69 μM artemisinin, respectively; *A. afra* PAR and LUX contained no detectable artemisinin. For *A. annua* tea infusion, an appropriate volume for the experimental design was added to the culture to yield a final artemisinin concentration of 7.78 μM. The same volume of the *A. afra* tea infusions was added to parasite cultures so that equivalent amounts of dry plant material were delivered for each infusion. This amounted to a 0.019 μM dose of artemisinin for SEN *A. afra* tea infusions, and undetectable artemisinin content in PAR and LUX *A. afra* tea infusions.

### *Plasmodium falciparum in vitro* culture

NF54 *P. falciparum* asexual parasites (gift of Dr. Ashley Vaughan, Seattle Children’s Research Institute) were maintained using standard conditions: a 37°C incubator at 5% O_2_ and 5% CO_2_ and maintained at 4% hematocrit using type A human erythrocytes (Red Cross) in complete media (CM: RPMI-1640 media supplemented with HEPES, D-glucose, hypoxanthine, gentamicin, sodium bicarbonate, and 10% type AB heat-inactivated human serum) (Moll *et al.*, 2013). Cultures were fed with daily media changes and diluted to 1% parasitemia every other day with fresh erythrocytes.

### Parasite synchronization and gametocyte formation protocol

Once asexual parasite cultures achieved 1% parasitemia or higher, they were synchronized for use in drug exposure assays. Working stock cultures were layered atop 70% Percoll columns (Percoll diluted in 10x RPMI, 13.3% sorbitol, and 1x PBS) and centrifuged at 2,500 x g for 10 min with no brake to yield a column with 4 distinct bands. The top two bands (media and infected erythrocytes) were removed from the column. Infected erythrocytes were washed with repeated cycles of adding incomplete media (ICM) (RPMI-1640 media supplemented with HEPES, D-glucose, hypoxanthine, gentamicin, and sodium bicarbonate), centrifugation, and removal of culture media. Synchronized erythrocytes were resuspended in CM to 2% or 4% hematocrit as dictated by the experimental design. To yield a gametocyte-rich culture for use in the various assays, we adapted a synchronization method from section 3.2 of Saliba and Jacobs-Lorena (Saliba and Jacobs-Lorena, 2013) and instead performed the above Percoll synchronization. After Percoll synchronization, culture medium was changed daily, erythrocytes were not replenished, and parasitemia was monitored by Giemsa stain. Once the culture reached 6-10% parasitemia, the CM was replaced by CM supplemented with 50 mM N-acetylglucosamine (NAG). NAG was used to eliminate asexual parasites and ensure an enriched gametocyte culture for testing. Daily medium changes were done with NAG-supplemented CM (NAG-CM) anywhere from 3-10 d depending on experimental design.

### Experimental design for drug exposure

For the asexual parasite drug exposure time course (Figure 1A), an asexual feeder culture was maintained using SCM. Cultures were synchronized using the aforementioned Percoll method and then the synchronized infected erythrocytes were split evenly into the appropriate number of T12.5 flasks as dictated by each experiment. Fresh erythrocytes, CM, and drug solution were added into each flask for a final hematocrit of 2%. Cultures were incubated in their respective drug treatments for 48 hr at standard conditions. For the early stage gametocyte drug exposure time course (Figure 1B), an asexual feeder culture was maintained according to SCM. Once asexual parasitemia reached at least 1%, the culture was Percoll synchronized and gametocyte formation was NAG induced. Cultures were treated with NAG-CM for 3 d after reaching the 6-10% parasitemia threshold. Prior to drug treatment, early stage gametocytes were washed with ICM, suspended in CM, and equal volumes of the suspension were aliquoted evenly into as many flasks as needed for the experimental design. Flasks were treated with drug and resuspended to 2% hematocrit. Cultures were incubated for 48 hr in standard culturing conditions. For the late stage gametocyte drug exposure time course (Figure 1C), the same protocol was followed as described for the early stage gametocyte assay, except cultures were maintained in NAG-CM for 12 d post-induction rather than 3 days post-induction. Artemisinin controls were prepared to a final concentration of 7.78 μM (high) or 0.019 μM (low) in 0.00275% DMSO. Methylene blue (MB) was prepared in water at a final concentration of 10 μM.

**Figure 1.**
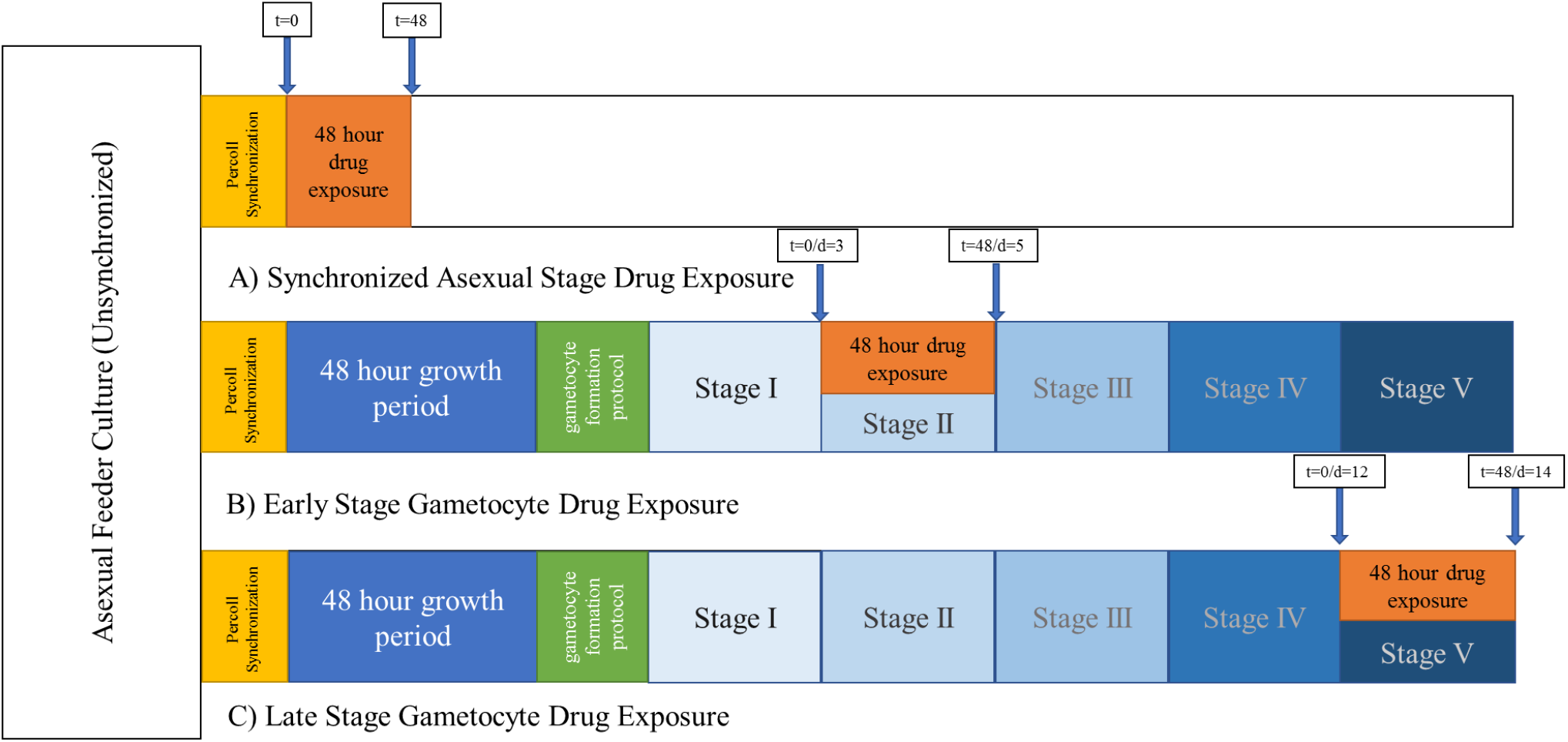
Experimental designs for each of the three assays used in this study. A) Timeline of asexual stage killing assay. B) Timeline of early stage gametocyte elimination assay. C) Timeline of late stage gametocyte elimination assay. Boxes with arrows indicate when samples were taken for RNA and or microscopy analysis.

### Microscopy analysis and morphology assessment

Thin-film smears were fixed in 100% methanol and Giemsa stained and counted using standard protocols (Moll *et al.*, 2013). Asexual parasitemia was determined and categorized as rings, trophozoites, or schizonts (Moll *et al.*, 2013). For absolute gametocyte counts, a 0.5 cm x 0.5 cm square was drawn on a thin-film Giemsa-stained smear and gametocytes were counted under 1000x magnification in that marked region. Erythrocytes were also counted in order to quantify gametocytemia. Each gametocyte was imaged and qualitatively assessed for morphological damage in order to score gametocyte ‘health’. ‘Healthy’ gametocytes had smooth, intact edges, a robust appearance, and possessed hemozoin crystals stained darker than the rest of the cell. ‘Unhealthy’ gametocytes had a sickly appearance characterized by a variety of cell membrane deformities (see examples shown above Table 2).

**Table 1.**
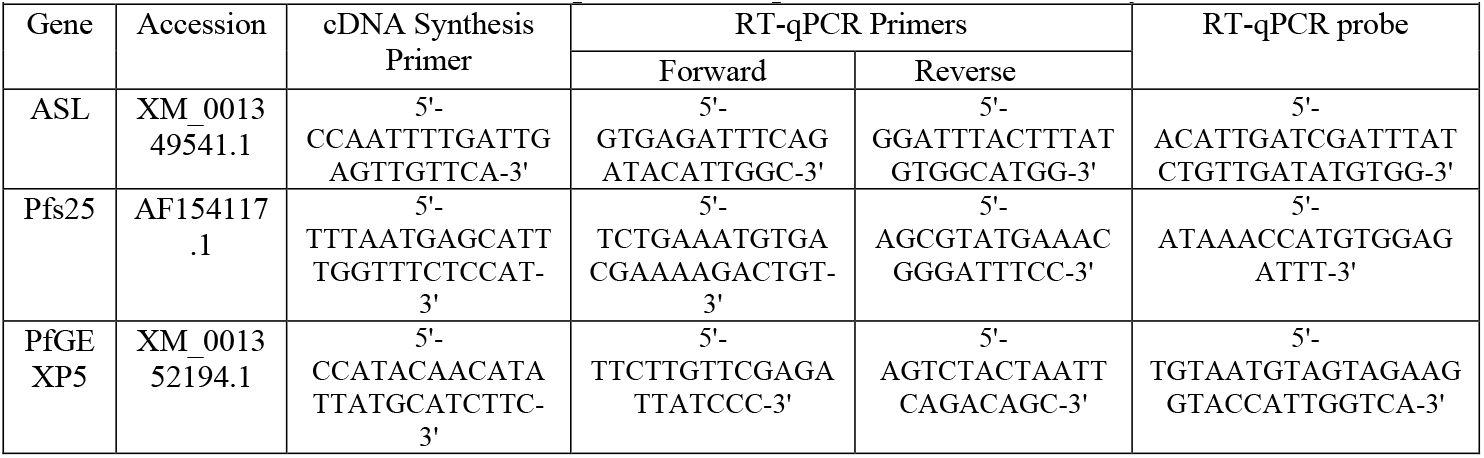
Genes, primers, and probes used in this study.

**Table 2.** Morphological aberrations among gametocytes before and after drug treatment.

| Treatment | Time (hr) | Total Gametocytes<sub>a</sub> | # Healthy Gametocytes | # Unhealthy Gametocytes | % Abnormal Morphology<sub>b,c</sub> | % Pinched Edges<sub>b,c</sub> | % Blebbed Edges<sub>b,c</sub> | % Bent<sub>b,c</sub> | % Jagged Edges<sub>b,c</sub> | % Lysed<sub>b,c</sub> | % Emaciated<sub>b,c</sub> |
| --- | --- | --- | --- | --- | --- | --- | --- | --- | --- | --- | --- |
| H<sub>2</sub>O control | 0 | 82 | 53 | 29 | 33.7 | 8.3 | 43.9 | 21.8 | 12.5 | 15.1 | 2.6 |
|  | 48 | 102 | 63 | 39 | 22.7 | 28.7 | 40.3 | 29.7 | 4.0 | 16.7 | 7.3 |
| DMSO control | 0 | 86 | 63 | 23 | 34.2 | 10.0 | 53.3 | 22.5 | 6.7 | 7.5 | 6.7 |
|  | 48 | 104 | 62 | 42 | 26.9 | 19.7 | 33.7 | 21.0 | 6.1 | 17.3 | 9.1 |
| Artemisinin (AN) | 0 | 58 | 44 | 14 | 22.5 | 12.5 | 55.0 | 27.5 | 12.5 | 5.0 | 0.0 |
|  | 48 | 82 | 45 | 37 | 17.4 | 9.1 | 46.6 | 8.0 | 7.3 | 20.2 | 14.3 |
| Methylene blue (MB) | 0 | 54 | 41 | 13 | 9.1 | 0.0 | 61.4 | 4.5 | 29.5 | 4.5 | 0.0 |
|  | 48 | 97 | 39 | 58 | 46.1 | 12.1 | 47.3 | 20.1 | 14.4 | 31.5 | 40.2 |
| SAM A. *annua* tea | 0 | 97 | 60 | 37 | 33.4 | 3.8 | 65.9 | 13.3 | 8.3 | 20.2 | 1.8 |
|  | 48 | 102 | 44 | 58 | 49.7 | 12.2 | 40.1 | 18.4 | 5.9 | 27.7 | 11.5 |
| SEN A. *afra* tea | 0 | 78 | 50 | 28 | 10.6 | 18.2 | 37.2 | 26.1 | 12.3 | 25.0 | 3.9 |
|  | 48 | 79 | 55 | 24 | 49.9 | 13.2 | 67.8 | 28.5 | 8.5 | 8.3 | 19.9 |

a: Sum of stage V gametocytes counted across three replicates for the time point and experimental condition.
b: Reflects percentage of abnormal gametocytes with specific deformity in that time point and treatment condition.
c: Total percentages may not add up to 100% because each percentage is the average of three replicates, and one single gametocyte may possess multiple forms of damage.

### RT-qPCR analysis of gametocyte-specific genes

Culture samples for RNA analysis were preserved using RNA*later* (Invitrogen) and stored at −20C until extraction. For extraction, RNA*later* reagent was removed, and then buffer ATL, Proteinase K, and buffer AL were added in that order from a QIAGEN QIAamp DNA mini kit. Samples were then processed using QIAGEN RNeasy mini kit, with the addition β-mercaptoethanol to RLT buffer and on-column DNA digestion using the QIAGEN RNase-free DNase kit. RNA was eluted in RNase-free water and stored at −80°C until use. cDNA was synthesized using QIAGEN QuantiTect Reverse Transcription kit using gene-specific primers (Table 1) and stored at −20°C until use. RT-qPCR reaction was performed in a Roche Lightcycler using FastStart Essential DNA Probes Master mix (Roche) and gene specific primer/probe sets (Table 1) with ASL as the reference gene.

### Reagents and other materials

All reagents were from Sigma-Aldrich unless otherwise already specified.

### Statistical analysis

Descriptive statistics of RT-qPCR were calculated using Excel. RT-qPCR data were analyzed using Excel and the Pfaffl method for determining relative gene expression (Pfaffl, 2001). Excel was also used for descriptive statistics on microscopy data. Data and statistical tests of RT-qPCR and microscopy data were analyzed using GraphPad Prism version 7.03. Normality of each dataset was determined using the Shapiro-Wilk normality test. Appropriate parametric or nonparametric tests were applied to the data sets based on the targeted comparison. Two-tailed paired *t-*test (or nonparametric equivalent) was used to compare time points within treatment conditions, whereas one-way ANOVA (or nonparametric equivalent) was used to compare between treatment conditions at a defined time point.

## Results

### *In vitro* assays show asexual stage elimination by *Artemisia* tea infusions

Prior to drug treatment, synchronized asexual cultures at 1% parasitemia consisted of approximately 72% trophozoites with no significant differences in culture composition between each treatment group (Figure 2).

**Figure 2.**
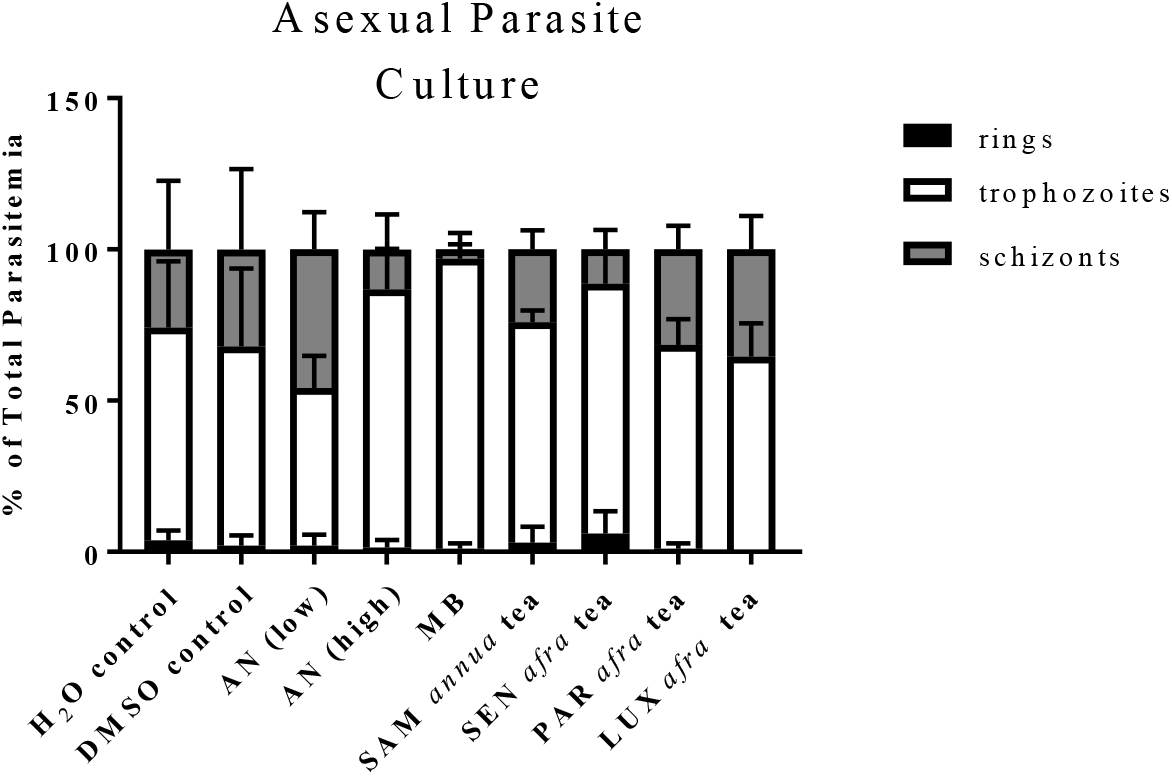
Asexual culture composition prior to drug treatment. Percent parasitemia of each asexual parasite stage was normalized to total parasitemia to determine proportions of life stages in each culture prior to drug treatment. AN, artemisinin; MB, methylene blue. Error bars = ± SD; *n* = 3. Kruskal-Wallis with Dunn’s multiple comparisons test performed on each parasite stage data set.

Thin blood smears were taken immediately after addition of drug and 48 hr after treatment. Asexual parasitemia increased in the H2O and DMSO-only controls over the 48 hr, showing uninhibited growth of asexual parasites during that time (Figure 3A). Parasitemia significantly decreased in the artemisinin and SAM *A. annua* and SEN *A. afra* infusion treatments, but not for the PAR and LUX *A. afra* tea infusion treatments (Figure 3A). Although SEN *A. afra* tea infusion yielded a significant level of inhibitory activity when compared to the H2O control at 48 hr (*p*=0.006), the inhibition was not as strong as that observed for the SAM *A. annua* tea infusion treatment (*p*<0.001) (Figure 3A). While both PAR and LUX *A. afra* tea infusions showed apparently similar growth inhibition to the SEN *afra* tea infusion, the results were not quite significant *(p* = 0.085 and 0.066, respectively). When the three *A. afra* cultivar tea infusions were compared to the SAM *A. annua* infusion, none of the *A. afra* infusions was significantly different (Figure 3B). This suggested that the three *A. afra* cultivar infusions have some inhibitory activity against *P. falciparum* parasites, but on a DW basis they were not as effective as SAM *A. annua* tea infusions. With confirmation that the experimental system was active against the asexual stage of *P. falciparum* parasites, we next determined how these two *Artemisia* tea infusions affected both early and late stage gametocytes.

**Figure 3.**
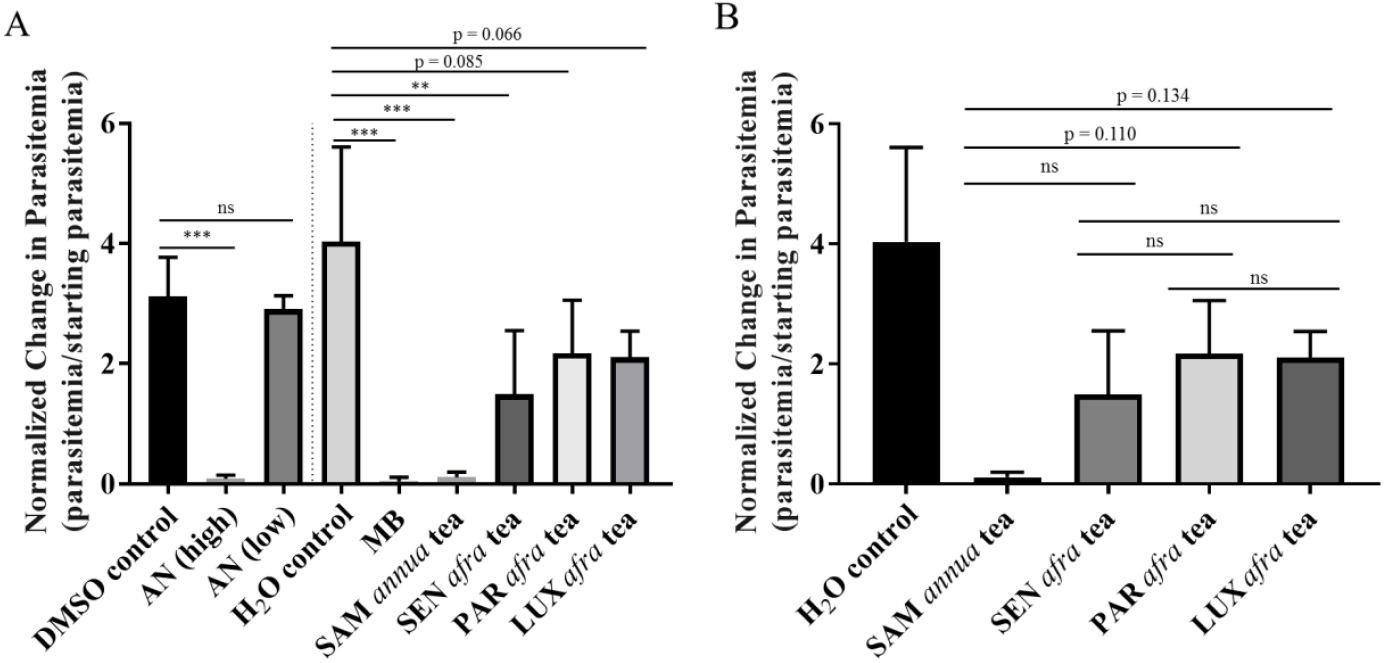
Microscopic determination of asexual parasitemia before and after drug treatment. A). Comparison of parasitemia after 48-hr treatment. B). Comparison of parasitemia after treatment with different *Artemisia spp.* tea infusions. Error bars, ±SD; *n* = 3, one-way ANOVA with Tukey’s multiple comparisons test, ns, not significant (*p*>0.05), ** = *p*≤0.01, *** = *p*≤0.001. AN, artemisinin; MB, methylene blue.

### Microscopically, *Artemisia* tea infusions eliminated *P. falciparum* gametocytes

*Artemisia* tea infusions were tested separately against early and late stage gametocytes. Stage III gametocytes are the earliest gametocyte stage that can be microscopically identified, so they were used as a proxy for the presence of stage I-III gametocytes. Prior to treatment, 55% of gametocytes counted were stage III gametocytes, with no significant differences between the individual cultures (Figure 4A).

**Figure 4.**
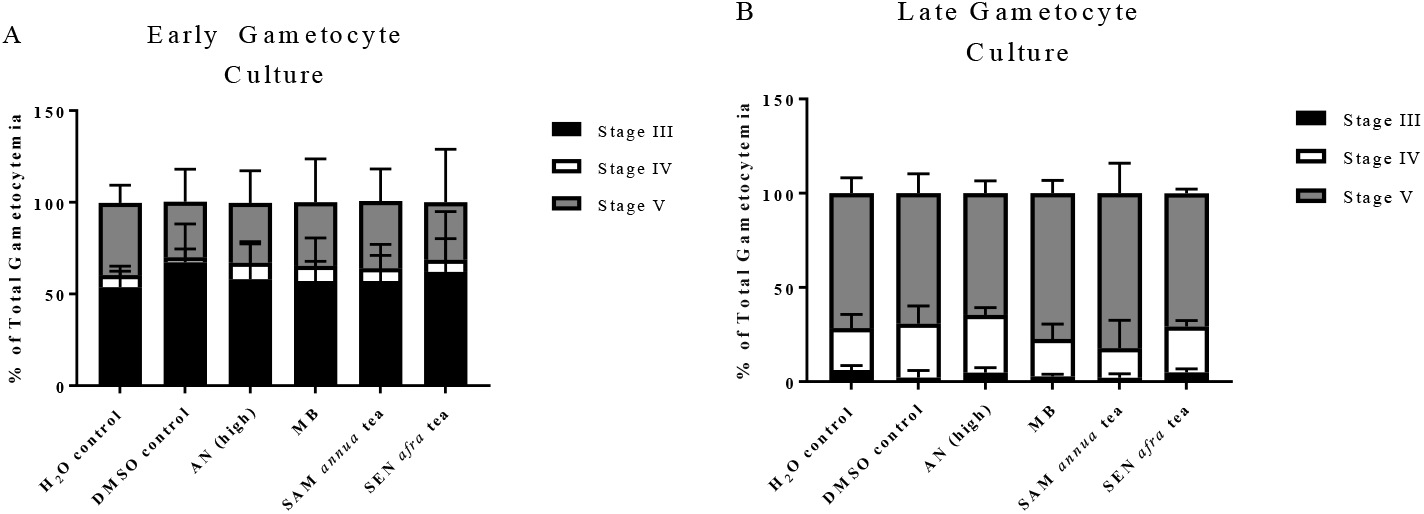
Gametocyte culture composition prior to drug treatment. A) Population composition of early gametocyte culture. B) Population composition of late gametocyte culture. Both culture compositions were determined by light microscopy. Percent gametocytemia of each gametocyte stage was normalized to total gametocytemia to determine proportions of life stages in each culture prior to drug treatment. AN, artemisinin; MB, methylene blue. Error bars = ± SD; *n* = 3. Kruskal-Wallis with Dunn’s multiple comparisons test performed on each parasite stage data set for late gametocyte culture, one-way ANOVA with Tukey’s multiple comparisons test performed on each parasite stage data set for early gametocyte culture.

After 48 hr treatment, stage III gametocytemia decreased significantly in the artemisinin and SAM *A. annua* tea infusion treatment groups, but there was no reduction in the SEN *A. afra* tea treatment group (Figure 5A).

**Figure 5.**
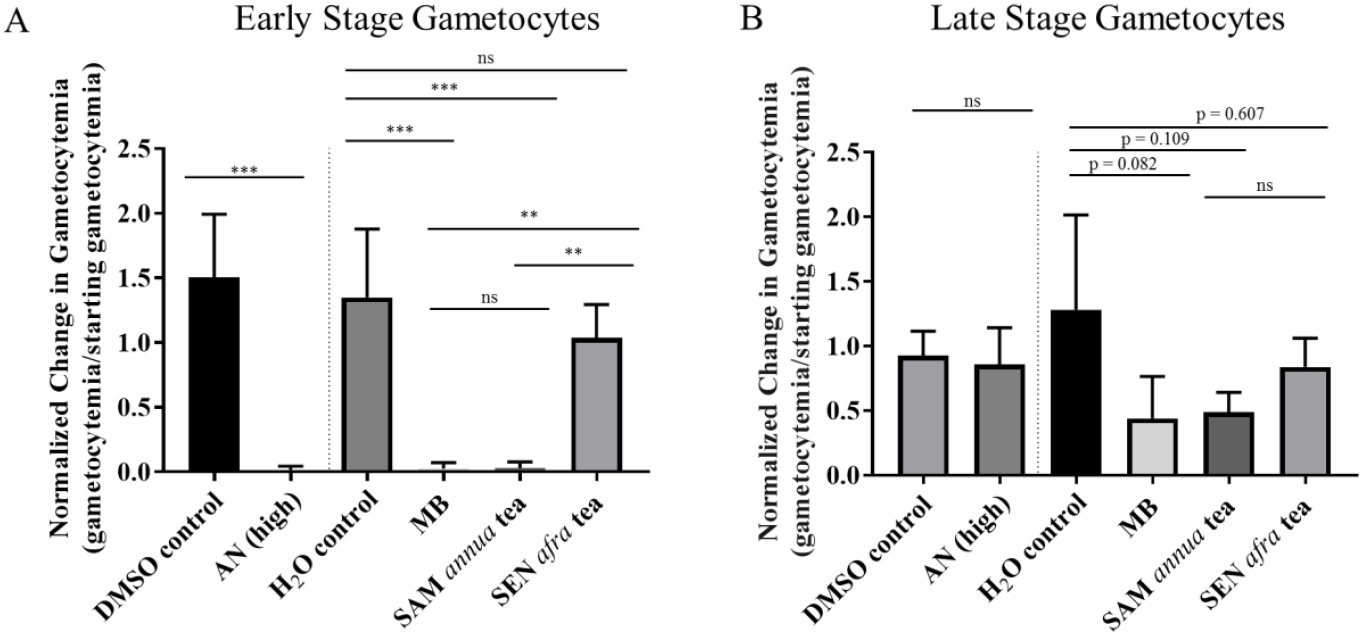
Healthy gametocytemia after 48-hour drug treatment as determined by light microscopy. A) Comparison of healthy stage III gametocytemia after 48-hr treatment. B) Comparison of healthy stage V gametocytemia after 48-hr treatment. Error bars, ± SD, *n* =3, one-way ANOVA with Tukey’s multiple comparisons test; ns, not significant (*p*>0.05), **= *p*≤0.01, ***=*p*≤0.001.

Prior to treatment, gametocyte cultures consisted of 73% healthy stage V gametocytes, with no significant difference between planned treatment conditions (Figure 4B). After 48 hr there were slight decreases in healthy stage V gametocytes after treatment with MB, and SAM *A. annua* and *A. afra* tea infusions (p = 0.082, p = 0.109, and p = 0.607, respectively), but all were insignificant compared to water controls (Figure 5B). These results were likely due to low overall gametocyte populations.

### Gametocyte morphology post-treatment reveals tractable and distinct types of damage

Besides counting gametocytes in cultures, the morphology of each gametocyte was scored to assess the overall health of that individual gametocyte. Healthy stage V gametocytes have a distinct, sausage-like shape with smooth, intact, and rounded edges; they appear plump (Table 2). Gametocytes were deemed unhealthy if they appeared emaciated, had bent or jagged edges, had abnormal bulging, or were lysed open (Table 2). Although this analysis depends on the assumption that only viable gametocytes can maintain a normal morphology, it provides additional information regarding how different treatments affected late stage gametocyte morphology. These damage data are summarized in Table 2 with representative images of observed morphologies illustrated along the top of Table 2. Although no significant differences could be found between treatments, there was generally more damage seen after 48 hr of treatment with MB and SAM *A. annua* tea infusions than with SEN *A. afra* tea infusions (Table 2).

### Artemisinin-containing treatments alter expression levels of gametocyte-specific genes

To probe more in depth, two different gametocyte-specific genes were measured using RT-qPCR, *PfGEXP5* and *Pfs25.* In early stage gametocytes, *PfGEXP5* expression was significantly reduced in SAM *A. annua* tea infusion treated cultures, and similar to the significant reduction observed in the pure artemisinin treated cultures. No effect was seen in the SEN *A. afra* treated cultures, suggesting that artemisinin content was a major driver of this effect (Figure 6A).

**Figure 6.**
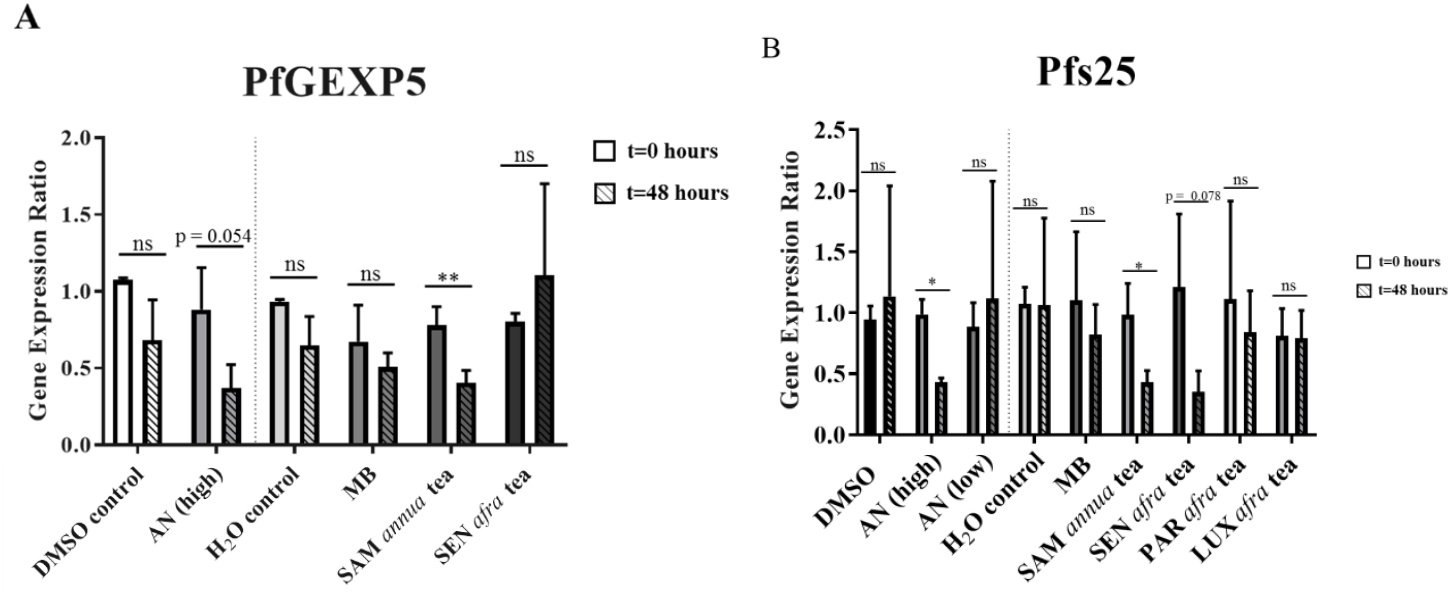
Quantification of gametocyte gene expression ratios via RT-qPCR in early and late stage gametocytes. A) *PfGEXP5* gene expression ratios before and after 48-hr treatment in early stage gametocytes. B) *Pfs25* gene expression ratios before and after 48-hr drug treatment in late stage gametocytes. Error bars, ± SD, *n*=3, two-tailed paired *t*-test (except for H_2_O control in B analyzed by Wilcoxon test), ns, not significant (*p*>0.05), *=*p*≤0.05. **=*p*≤0.01.

A similar effect was seen in late stage gametocytes for *Pfs25.* There were significant reductions in expression in cultures treated with pure artemisinin (7.78μM, but not seen at 0.019 μM) and SAM *A. annua* tea infusion treatment groups. There was also a nearly significant (p=0.078) reduction for the SEN *A. afra* tea infusion treatment condition (Figure 6b). Interestingly, despite MB’s gametocytocidal activity seen in the microscopy results, MB did not reduce the expression of either gametocyte-specific gene tested here. Taken together, these results suggest that there may be an artemisinin-specific effect on these two gametocyte-specific genes.

## Discussion

To our knowledge, this is the first study that measured the *in vitro* antiparasitic ability of *Artemisia* tea infusions against the sexual, gametocyte stages of *P. falciparum*. Several interesting and relevant patterns have emerged. First, there appears to be a correlation between artemisinin concentration and antiparasitic efficacy of the tea infusions *in vitro*. Duffy and Avery (2013) used parasite strain NF54 to determine the IC_50_ of pure artemisinin against early and late stage gametocytes, and the IC_50_s were 12 nM and 5 nM, respectively. Although tea infusions contain other phytochemicals besides artemisinin, they still behaved in a dosedependent manner. According to the published IC_50_s, both SAM *A. annua* and SEN *A. afra* tea infusions delivered enough artemisinin to kill late stage gametocytes, but only SAM *A. annua* delivered enough artemisinin to eliminate early stage gametocytes, results consistent with the published IC_50_ (Duffy and Avery, 2013). For early stage gametocytes the results of this study were fully consistent with the Duffy and Avery (2013) results; there was a significant decrease in gametocytemia when exposed to SAM *A. annua* tea infusions, but not SEN infusions. For late stage gametocytes, both SAM and SEN had anti-gametocyte activity, but not as powerful as anticipated. This was attributed to the fact that in the late stage experiments the gametocyte populations were lower than those measured at the early stages closer to NAG induction. Results of this study, thus, were generally consistent with the results of Duffy and Avery (2013) and showed better efficacy for the SAM vs. the SEN infusions. Taken together, these suggested that artemisinin was the major driver of gametocytocidal activity for these *Artemisia* tea infusions.

Although at first glance these results appear inconsistent with the clinical trial data of Munyangi et al (2019) that showed both *A. afra* and *A. annua* tea infusions eliminated gametocytes, they are consistent when considering the total delivered amount of artemisinin. In this study 7.78 μM artemisinin was delivered from the SAM *A. annua* tea, whereas only 19 nM artemisinin was delivered from the SEN *A. afra* tea. However, in the clinical trial, each patient received a minimum of 520 nM artemisinin from the *A. afra* tea infusion per dose, and the *A. annua* tea infusion delivered over 45 times the amount delivered in the *A. afra* tea infusion (Munyangi *et al.*, 2019). The clinical trial showed that *A. afra* tea infusion was an equally viable treatment option against *P. falciparum* malaria despite its negligible artemisinin content. The *in vitro* results of this study showed that artemisinin content was the main factor in assessing the *in vitro* gametocidal efficacy of these teas.

Along with microscopically quantifying gametocytemia after treatment, we were also able to score the health of gametocytes via morphological analysis. Although we were unable to determine any significant differences between treatment groups after 48 hr treatment, in general there was a higher percentage of damaged gametocytes in cultures treated with MB or SAM *A. annua* tea infusion. The results obtained by using MB were consistent with Wadi et al. (2018) who showed that MB is effective at eliminating both early and late stage gametocytes, with an IC_50_ of 424 nM and 106 nM, respectively. They also showed that MB induced distinct morphological damage to late stage gametocytes, including membrane deformities or shrinkage (Wadi *et al.*, 2018). After 48 hr treatment with 10 μM MB, we also observed substantial morphological damage in the form of various membrane deformities. This suggests that the morphological damage observed in the SAM *A. annua* tea infusion may indicate that the additional phytochemicals present in the infusion contributed to the overall antiparasitic effect, particularly because similar levels of damage were not seen in the artemisinin-only treatment condition.

We were also interested in exploring what was occurring on a molecular level when gametocytes were treated with *Artemisia spp.* tea infusions. To address this question, we looked at two gametocyte-specific genes. *PfGEXP5* is the earliest known gametocyte-specific gene to be expressed (Tibúrcio *et al.*, 2015), and *Pfs25* is a stage V gametocyte marker gene expressed predominantly in late stage gametocytes. *Pfs25* function is well characterized as an ookinete surface antigen that is translationally repressed in the late stage female gametocyte (Kaslow *et al.*, 1988). *PfGEXP5* function is currently unknown, although it is expressed about 14 hours after a sexually committed merozoite invades an erythrocyte, and that it is likely exported into the host cell cytoplasm to perform its function (Tibúrcio *et al.*, 2015).

Here we showed that *PfGEXP5* and *Pfs25* expression levels decreased when there were appreciable amounts of artemisinin in the treatments, but not so in the MB treatment. Nevertheless, MB significantly reduced microscopic counts of gametocytes. This suggests that the gametocytocidal effects of each treatment was due to distinctly different mechanisms of action leading to different gametocyte-specific gene expression profiles. Although the mechanism of action for both artemisinin and MB are not fully elucidated, it is thought that MB is an oxidative stress inducer that specifically targets the cellular antioxidant protein glutathione reductase (Delves *et al.*, 2013; Mott *et al.*, 2015). Artemisinin likely has multiple mechanisms of action. When the molecule comes into contact with free heme in the parasite, the endoperoxide bridge is cleaved and reactive oxygen species (ROS) are released, causing damage to parasite proteins via alkylation (Medhi *et al.*, 2009; Delves *et al.*, 2013) The molecule itself can also bind directly to at least 124 different parasite proteins (Wang *et al.*, 2015). Since neither *Pfs25* nor *PfGEXP5* play a role in the oxidative stress response, it follows that neither of these genes are targets of MB. However, since artemisinin targets at least 124 proteins from all different cellular processes, it is possible that both *PfGEXP5* and *Pfs25* are direct targets of artemisinin.

Although the functional importance of *PfGEXP5* has not yet been elucidated, *Pfs25* yields the ookinete surface antigen, and reducing the expression of this gene may have downstream effects in the female gamete. In fact, there is evidence that P25 is essential for ookinete survival in the mosquito midgut, as well as transformation into an oocyst (Tomas *et al.*, 2001). Further research is needed to understand the functional relevance of this decrease in expression of gametocyte-specific genes.

In this study, we aimed to determine the antiparasitic effects of various *Artemisia* tea infusions on different stages of *P. falciparum* gametocytes. At the time of this report, there were at least seven other studies that tested *Artemisia* tea infusions against *P. falciparum in vitro* (Liu *et al.*, 2010; de Donno *et al.*, 2012; Silva *et al.*, 2012; Mouton *et al.*, 2013; Omar *et al.*, 2013; Suberu *et al.*, 2013; Zime-Diawara *et al.*, 2015). However, all of these studies only looked at asexual parasites. The asexual parasite data of this study aligns well with results published in those studies. Of the seven, two reported IC_50_s in nM, with an IC_50_ of 2.9-7.6 nM (Suberu *et al.*, 2013; Zime-Diawara *et al.*, 2015). The SEN *A. afra* tea infusion delivered 19 nM artemisinin per dose, and the SAM *A. annua* tea infusion delivered about 400x that amount (7.6 μM). The minimum threshold of artemisinin for killing *P. falciparum* asexual parasites is reported at ~10 μg/L (0.035 μM) (Alin and Bjorkman, 1994), so while the SAM infusion was >200 fold greater than the minimum artemisinin level, the SEN infusion had about half the artemisinin concentration required to kill asexual parasites and yet there was a significant decrease in parasitemia within 48 hr. Despite having undetectable levels of artemisinin, *A. afra* PAR and LUX tea infusions also decreased asexual parasitemia comparable to that observed with SEN. Together these results suggest that there are other synergistic or antimalarial compounds present in these *Artemisia* cultivars that are providing this antiparasitic activity despite the absence of detectable artemisinin.

Furthermore, when the antiparasitic efficacy of the low artemisinin control treatment is compared to the efficacy of SEN *A. afra* tea infusion, both of which delivered 19 nM artemisinin, there appeared to be a stronger effect due to the tea infusion than to the pure artemisinin (although it was not a significant difference). This is consistent with other reports where an IC_50_ of AN delivered by *Artemisia* tea infusions against *in vitro* asexual *P. falciparum* parasites was 2.9-7.6 nM (Suberu *et al.*, 2013; Zime-Diawara *et al.*, 2015), whereas the IC_50_ of pure artemisinin against *in vitro* asexual *P. falciparum* parasites was 42 nM (Duffy and Avery, 2013). In contrast to the recent claim by Czechowski, et al. (2019), these results support the hypothesis that *Artemisia* tea infusions are antiparasitic and deliver additional phytochemicals that likely act either synergistically with artemisinin to enhance its antimalarial ability or possess their own antimalarial activity.

## Conclusions

This study provides *in vitro* evidence that *Artemisia spp.* tea infusions had gametocytocidal activity against both early and late stage gametocytes, but with differential effects in gametocidal activity, in morphological aberrations, and in gene expression. *Artemisia* tea infusions that contain little to no artemisinin were also antiparasitic, but less so than cultivars containing artemisinin. *Artemisia* tea infusions are also effective against the asexual stages of the parasite but become less effective as the artemisinin content of a specific cultivar declines. No *in vitro* study using any extract can replicate *in vivo* studies in which there are also likely host interactions with the therapeutic. Nevertheless, this study is consistent with the prior human malaria clinical trials showing that *Artemisia* tea infusions are gametocidal. Further work is needed to determine if the tea infusions also prevent transmission to the mosquito vector. If that occurs, then *Artemisia spp.,* especially those containing reasonable amounts of artemisinin (~1%), could provide a more cost-effective means to thwart this deadly disease.

## Acknowledgements

Thank you to Dr. Ashley Vaughan (Seattle Children’s Research Institute) for the NF54 parasite line, Dr. Lisa Checkley (University of Notre Dame) for assistance with troubleshooting parasite synchronization, to Dr. Ann Stewart’s lab, Uniformed Services University of Health Sciences, Bethesda, MD for training DS in efficiently extracting parasite RNA from blood samples, and to Dr. Melissa Towler for plant sample analyses and parasite counting. SEN, PAR, and LUX were from Dr. Guy Mergei, University of Liege, Belgium, Dr. Lucille Cornet-Vernet from La Maison d’Artemisia, France, and Dr. Pierre Lutgen, Luxemborg, respectively. This work was funded by the Ararat fund for Artemisinin Research at WPI. Phytochemical analysis of *Artemisia* samples was supported in part by The National Center for Complementary and Integrative Health, award number NIH-2R15AT008277-02. The content is solely the responsibility of the authors and does not necessarily represent the official views of the National Center for Complementary and Integrative Health or the National Institutes of Health.

## Author contributions

DS designed experiments, conducted experiments, analyzed data, wrote manuscript. PW designed experiments, analyzed data, wrote manuscript.

